# No supporting evidence of classification based on FFPE samples, ambiguity in classification of EGFR mutants, and inclusion of bona-fide platelet genes in discriminator sets indicate no biological basis for using RNA-seq from tumor-educated platelets as a source in ”liquid biopsy”

**DOI:** 10.1101/146134

**Authors:** Sandeep Chakraborty

## Abstract

In this detailed critique of the study proposing using RNA-seq from tumor-educated platelets (TEP) as a ‘liquid biopsy’ source [1], several flawed assumptions leave little biological basis behind the statistical computations. First, there is no supporting evidence provided for the FFPE based classification of MET-overexpression and EGFR mutation on tumor-tissues. Considering that raw reads of MET expression in a subset of healthy [N=21, mean=112, sd=77] and NSCLC [N=24, mean=11, sd=12] samples (typically with millions of reads) translates into over-expression in reality, providing the data for such computations is vital for future validation. A similar criticism applies for classifying samples based on EGFR mutations (the study uses only exon 20 and 21 from a wide range of possible mutations) with negligible counts [N=24, mean=3, sd=6]. While Ofner et. al, 2017 faced ‘major problems associated with FFPE DNA’, it is also true that Fassunke, et al., 2015 found concordance in 26 out of 26 samples for EGFR mutations in another FFPE-based study. However, Fassunke, et al., 2015 have been meticulous in describing the EGFR amplicons (exon 18 and 19 are missing in the TEP-study). Any error in initial classification renders downstream computations error-prone. The low counts of MET in the RNA-seq firmly establishes that inclusion of genes with such low counts in the set of 1100 discriminatory genes (Table S4) makes no sense as the “real” counts could vary wildly. Yet, TRAT1 is an example of one discriminator gene with counts of healthy [N=21, mean=164, sd=375] and NSCLC [N=24, mean=53, sd=176]. There are many such genes which should be excluded. Moving on to a discriminator with high counts (F13A1) in both healthy [N=21, mean=28228, sd=48581] and NSCLC [N=24, mean=98336, sd=74574] samples, a bonafide platelet gene that “encodes the coagulation factor XIII A subunit”. Platelets do not have a nucleus, and thus the blue-print (chromosomes and related machinery) for making or regulating mRNA. They are boot-strapped with mRNA, like F13A1, during origination and then just go on keep collecting mRNA during circulation (which is the premise of their use in liquid biopsy). The assumption that these genes are differentially spliced in huge numbers is highly speculative without providing experimental proof. The discovery of spliceosomes in anucleate platelets [2] in 2005, 30 years after splicing was discovered in the nucleus by Sharp and Robert, probably indicates that spliceosomes are not dominant in platelets. Zucker, et al., 2017 have shown for another gene F11 that it ‘is present in platelets as pre-mRNA and is spliced upon platelet activation’ [3]. Any study using the F13A1 gene as a discriminator ought to show the same two things, followed by differential counts in TEP. Ironically, F11 is not present in the discriminator set. Another blood coagulation related gene (TFPI) shows slight over-expression in NSCLC (moderate counts, healthy [N=21, mean=1352, sd=592] and NSCLC [N=24, mean=1854, sd=846]), agreeing with Iversen, et al., 1998 [4], but in contrast to Fei, et al., 2017 [5], demonstrating that the jury is still out on the levels of many such genes. Thus, circulating mRNA from tumor tissues are not discriminatoryif MET is degraded to such levels in platelets ‘educated’ by NSCLC tumors, why not other possible mRNA that might have been picked during the same ‘class’? Furthermore, high count genes can only be bona-fide platelet genes, and have no supporting experimental proof of splicing differences (any one gene would suffice to instill some confidence). In conclusion, looking past the statistical smoke surrounding “surrogate signatures”, one finds no biological relevance.

## Introduction

Tumor tissue biopsy, the gold standard for cancer diagnostics, pose challenges that include access to the tumor, quantity and quality of tumoral material, lack of patient compliance, repeatability, and bias of sampling a specfic area of a single tumor [6]. This has resulted in a new medical and scientific paradigm defined by minimal invasiveness, high-efficiency, low-cost diagnostics [7], and, whenever possible, personalized treatment based on genetic and epigenetic composition [8]. The presence of fragmented DNA in the cell-free component of whole blood (cfDNA) [9], first reported in 1948 by Mandel and Metais, has been extensively researched for decades, with extremely promising results in certain niches [10]. Additionally, cfDNA derived from tumors (ctDNA) [11] have tremendous significance as a cancer diagnostic tool [12], and for monitoring responses to treatment [13]. However, detection of ctDNA, and differentiation with cfDNA, remains a challenge due the low amounts of ctDNA compared to cfDNA [14].

Recently, tumor-educated blood platelets (TEP) were proposed as an alternative source of tumor-related biological information [1, 15]. The hypothesis driving the potential diagnostic role of TEPs is based on the interaction between blood platelets and tumor cells, subsequently altering the RNA profile of platelets [16,17]. The study showed using RNA-seq data that tumor-educated platelets (TEP) can distinguish 228 patients with localized and metastasized tumors from 55 healthy individuals with 96% accuracy [1]. As validation, this study reported significant over-expression of MET genes in non-small cell lung carcinoma (NSCLC), and HER2/ERBB2 [18] genes in breast cancer, which are well-established biomarkers. Also, using a set of 1072 genes, the study reported *>*95% accuracy in pan-cancer diagnostics (Fig 1 in [1]).

Here, the TEP-study is refuted based on the absence of any kind of biological relevance to the discriminator set of genes other than varying numbers, which could be assigned to a number of reasons. First, the almost negligible amounts of MET/EGFR reads in the NSCLC samples is demonstrated. The critiques can be enumerated as:

1. There is no supporting evidence for FFPE based classification of MET-overexpression and EGFR mutants.
2. Inclusion of genes with such low counts, like TRAT1.
3. Genes with high reads, like F13A1, which can only be bona-fide platelet genes.

### Contradictory results

#### Erroneous and ambiguous classification of MET-overexpression and EGFR mutations

Both MET-overexpression and EGFR mutations use FFPE, which are solutions ‘designed to meet the challenges of analyzing degraded or limited genomic material’ (https://www.illumina.com/science/education/ffpesample-analysis.html). ‘Overexpression of MET protein in tumor tissue relative to adjacent normal tissues occurs in 25-75% of NSCLC and is associated with poor prognosis’ https://www.mycancergenome.org/content/disease/lung-cancer/met/343/. However, in the TEP-study only 13% (8 out of 60) are MET+, which seems too low (Table S1). Further, the classification “MET WT” is not clearly defined.

Similarly, for EGFR mutations (exon 20 and 21 spanning from 2541 to 2881) gives a small subset of possible EGFR mutations. A comprehensive list is provided in https://www.mycancergenome.org/content/disease/lungcancer/egfr/339/: Kinase Domain Duplication, c.2156G*>*C (G719A), c.2155G*>*T (G719C), c.2155G*>*A (G719S), Exon 19 Deletion, Exon 19 Insertion, Exon 20 Insertion, c.2290 2291ins (A763 Y764insFQEA), c.2303G*>*T (S768I), c.2369C*>*T (T790M), c.2573T*>*G (L858R) and c.2582T*>*A (L861Q).

While Ofner, et al., 2017 ‘were not able to confirm any assumed hotspot mutation within repeated sequencing of relevant amplicons suggesting the detection of sequence artifacts most likely caused by DNA lesions associated with FFPE tissues’ [19], Fassunke, et al., 2015 found concordance in 26 out of 26 samples for EGFR mutations, supporting FFPE studies [20]. However, they provide a complete list of the amplicons in Table 1, a meticulous classification of the EGFR mutations found in Table 4, and macrodissection showing EGFR mutation status in Fig 1 [20]. This includes EGFR exons 18 and 19 as well, which have been missed out in the TEP study (‘EGFR (exon 20 and 21) amplicon deep sequencing strategy ($5,0003 coverage) on the Illumina Miseq platform using prospectively collected blood samples of patients with localized or metastasized cancer’ [1]).

To summarize, for MET-overexpression classification (with an exceptionally low occurrence compared to known values) the FFPE data has not been provided. In the case of EGFR classification in the TEP-study, it clearly is erroneous, since only a subset of known mutations were used. It is vital to know that the classification has been done properly, before providing statistics based on surrogate biomarkers.

### Genes with such low reads, like TRAT1, can not be discriminators

Since the raw counts of healthy and NSCLC samples translate (and rightly in most cases), to over-expression of MET in NSCLC samples,

Healthy MET = [287 197 61 142 178 127 7 176 2 133 188 156 23 2 185 170 104 23 28 71 108]

NSCLC MET = [0 14 24 5 11 9 45 1 5 11 9 12 5 2 34 3 7 42 6 16 12 2 5 2],it is obvious that genes with small counts (say in the hundreds) can not be discriminators.

Yet, TRAT1 is one such gene in the set of 1072 genes that has been shown to give *>*95% accuracy as a pan-cancer diagnostic (Fig 1 in [1], Table S4). The counts of TRAT1 are as low as those of MET.

Healthy = [158 75 39 88 98 96 0 242 0 92 359 53 7 3 53 51 103 11 1801 48 67]

NSCLC = [0 1 8 0 40 1 855 0 8 10 2 25 1 288 19 0 0 3 1 3 2 0 13 0]

Though, it does show empirical statistical difference, it could be the exact opposite in reality (as with the MET gene).

### Genes with high counts, like F13A1, are bona-fide platelet genes, and can not be discriminator genes

F13A1, a bona fide platelet gene that “encodes the coagulation factor XIII A subunit”, in the set of 1072 discriminator has high counts:

Healthy = [21927 14452 17792 16123 8889 48109 17111 21839 21349 12212 50801 21673 1141 15708 7446 9451 6448 6446 238681 26216 8994]

NSCLC = [314955 17989 95737 115057 65335 39798 251593 131285 41424 92801 120105 90751 69381 124199 109380 229636 80741 3821 109441 24817 9621 51099 123293 47826]

Platelets have no nucleus, and are bootstrapped with their mRNA during origination [21]. Phillip Sharp and Richard Roberts discovered splicing in 1977, for which they were awarded the Nobel Prize in 1993. Spliceosomes were assumed to be nucleus-confined till 2005, when it was a surprising find that ‘primary human platelets also contain essential spliceosome factors including small nuclear RNAs, splicing proteins, and endogenous pre-mRNAs’‘[2]. The 30 year lag in this realization probably indicates that splicing can not be a dominant effect in platelets, and the causal agent of very large differences in counts. So, it would be speculative to implicate splicing to account for the difference in F13A1 counts, and assign it discriminatory status, without proper experimental proof. For example, Zucker, et al., 2017 have shown that ‘that F11 is present in platelets as pre-mRNA and is spliced upon platelet activation’ [3]. In addition to demonstrating the same for F13A1, any study using this gene as a discriminator ought to show differential counts in TEP. Ironically, F11 is not present in the discriminator set.

### The jury is out on the expression level of genes, like TFPI

The ambiguity hidden in the discriminator set of 1072 is shown through another gene related to blood coagulation TFPI. TFPI has the following counts:

Healthy = [1573 1573 562 1803 1174 1479 1729 1515 1137 1077 2119 1517 189 1801 1036 872 1709 206 2811 1171 1343]

NSCLC = [4027 3250 2118 1700 1290 1095 2908 1931 1558 3280 2207 2161 1837 1776 2150 2277 540 729 1444 1048 1316 694 1901 1268]

This shows slight over-expression in NSCLC, in agreement with Iversen, et al., 1998 [4], but in contrast to Fei, et al., 2017 [5], demonstrating that the jury is still out on the levels of many such genes. Decreasing TFPI levels in NSCLC makes more biological sense, since it leads to activation of blood coagulation, which contributes to cancer progression. Additionally, one might also argue about the read counts are moderate, possibly tending to very low levels, where they are not discriminatory.

## Conclusion

This study raises serious doubts on using TEP as a possible ‘liquid biopsy’ candidate. Essentially, it refutes the hypothesis that platelets carry enough RNA-seq from tumors to make it viable as a diagnostic method. This has been vaguely worded in the TEP-study ‘contained undetectable or low levels of these mutant biomarkers’ [1], suggesting that other mRNA (”surrogate signatures”) might encode enough information for cancer diagnostics. Here, it is shown in details that most of the 1072 discriminator genes make no sense. The onus lies on the authors of the study to show at least one gene that is differentially regulated in proximity to tumor cells to prove some sort of biological relevance.

Again the statement ‘Further validation is warranted to determine the potential of surrogate TEP profiles for blood-based companion diagnostic’ is a truism. Further validation is obviously needed what has been missed is a deeper look at the data presented in the current study. A review found it ‘surprising’ that although ‘the tumor type was the predominant factor for the actual platelet conditioning, tumor metastasis did not significantly impact on them when compared to samples from patients without metastasis’ [17]. The excitement surrounding the fact that ‘2016 marked the first approval of a liquid biopsy test in oncology to assist in patient selection for treatment’ [22] should be tempered, and a cautious approach adopted [23, 24] with reports of ‘broken promises’ [25].

Circulating mRNA from tumor tissues are not discriminatory if MET is degraded to such levels in platelets ‘educated’ by NSCLC tumors, why not other possible mRNA that might have been picked during the same ‘class’? High count genes can only be bona-fide platelet genes, and have no supporting proof of splicing differences (any one gene would suffice). In conclusion, looking past the statistical smoke surrounding “surrogate signatures”, one finds no biological relevance.

## Materials and methods

A smaller subset of lung cancer samples (DATASET:list.lung.txt,n=24) and healthy (DATASET:list.healthy.txt,n=21) was used from the given 60 NSCLC samples in the TEP study [1]. A kmer-based version (KEATS [26]) of YeATS [27–31] was used to obtain gene counts from transcripts in the RNA-seq data. The counts provided are raw reads, and not normalized - but does not alter the criticisms. A BLAST search suffices to demonstrate the absence of MET/EGFR genes in the lung cancer RNA-seq samples.

## References

1. Best MG, Sol N, Kooi I, Tannous J, Westerman BA, et al. (2015) Rna-seq of tumor-educated platelets enables blood-based pan-cancer, multiclass, and molecular pathway cancer diagnostics. Cancer cell 28: 666–676.

2. Denis MM, Tolley ND, Bunting M, Schwertz H, Jiang H, et al. (2005) Escaping the nuclear confines: signal-dependent pre-mrna splicing in anucleate platelets. Cell 122: 379–391.

3. Zucker M, Hauschner H, Seligsohn U, Rosenberg N (2017) Platelet factor xi: Intracellular localization and mrna splicing following platelet activation. Blood Cells, Molecules, and Diseases.

4. Iversen N, Lindahl AK, Abildgaard U (1998) Elevated tfpi in malignant disease: relation to cancer type and hypercoagulation. British journal of Haematology 102: 889–895.

5. Fei X, Wang H, Yuan W, Wo M, Jiang L (2017) Tissue factor pathway inhibitor-1 is a valuable marker for the prediction of deep venous thrombosis and tumor metastasis in patients with lung cancer. BioMed research international 2017.

6. Vendrell JA, Mau-Them FT, Béganton B, Godreuil S, Coopman P, et al. (2017) Circulating cell free tumor dna detection as a routine tool forlung cancer patient management. International Journal of Molecular Sciences 18: 264.

7. Han X, Wang J, Sun Y (2017) Circulating tumor dna as biomarkers for cancer detection. Genomics, proteomics & bioinformatics.

8. Sorber L, Zwaenepoel K, Deschoolmeester V, Van Schil P, Van Meerbeeck J, et al. (2016) Circulating cell-free nucleic acids and platelets as a liquid biopsy in the provision of personalized therapy for lung cancer patients. Lung Cancer.

9. Jiang P, Lo YD (2016) The long and short of circulating cell-free dna and the ins and outs of molecular diagnostics. Trends in Genetics 32: 360–371.

10. Lo YD, Corbetta N, Chamberlain PF, Rai V, Sargent IL, et al. (1997) Presence of fetal dna in maternal plasma and serum. The Lancet 350: 485–487.

11. Chen XQ, Stroun M, Magnenat JL, Nicod LP, Kurt AM, et al. (1996) Microsatellite alterations in plasma dna of small cell lung cancer patients. Nature medicine 2: 1033–1035.

12. Yi X, Ma J, Guan Y, Chen R, Yang L, et al. (2017) The feasibility of using mutation detection in ctdna to assess tumor dynamics. International Journal of Cancer 140: 2642–2647.

13. Imamura F, Uchida J, Kukita Y, Kumagai T, Nishino K, et al. (2016) Monitoring of treatment responses and clonal evolution of tumor cells by circulating tumor dna of heterogeneous mutant egfr genes in lung cancer. Lung Cancer 94: 68–73.

14. Diaz LA, Bardelli A (2014) Liquid biopsies: genotyping circulating tumor dna. Journal of Clinical Oncology 32: 579–586.

15. Nilsson RJA, Balaj L, Hulleman E, Van Rijn S, Pegtel DM, et al. (2011) Blood platelets contain tumor-derived rna biomarkers. Blood 118: 3680–3683.

16. Bardelli A, Pantel K (2017) Liquid biopsies, what we do not know (yet). Cancer cell 31: 172–179.

17. Feller SM, Lewitzky M (2016) Hunting for the ultimate liquid cancer biopsy-let the tep dance begin. Cell Communication and Signaling 14: 24.

18. Foulkes WD, Stefansson IM, Chappuis PO, Bégin LR, Goffin JR, et al. (2003) Germline brca1 mutations and a basal epithelial phenotype in breast cancer. Journal of the National Cancer Institute 95: 1482–1485.

19. Ofner R, Ritter C, Ugurel S, Cerroni L, Stiller M, et al. (2017) Non-reproducible sequence artifacts in ffpe tissue: an experience report. Journal of cancer research and clinical oncology.

20. Fassunke J, Haller F, Hebele S, Moskalev EA, Penzel R, et al. (2015) Utility of different massive parallel sequencing platforms for mutation profiling in clinical samples and identification of pitfalls using ffpe tissue. International journal of molecular medicine 36: 1233–1243.

21. Harrison P, Goodall AH (2008) message in the platelet–more than just vestigial mrna! Platelets 19: 395–404.

22. Blumenthal GM, Pazdur R (2017) Approvals in 2016: the march of the checkpoint inhibitors. Nature Reviews Clinical Oncology 14: 131–132.

23. Diamandis EP (2016) A word of caution on new and revolutionary diagnostic tests. Cancer cell 29: 141–142.

24. Best MG, Sol N, Tannous BA, Wesseling P, Wurdinger T (2016) Re: a word of caution on new and revolutionary diagnostic tests. Cancer cell 29: 143.

25. Shee K, Chamberlin M, Varn F, Bean J, Marotti J, et al. (2017). Abstract p6-07-03: Broken promise of liquid biopsy: Plasma dna does not accurately reflect tumor dna in metastatic breast cancer.

26. Chakraborty S (2017) Cataloguing over-expressed genes in epstein barr virus immortalized lymphoblastoid cell lines through consensus analysis of pacbio transcriptomes corroborates hypomethylation of chromosome 1. bioRxiv: 125823.

27. Chakraborty S, Britton M, Wegrzyn J, Butterfield T, Martinez-Garcia PJ, et al. (2015). YeATS-a tool suite for analyzing RNA-seq derived transcriptome identifies a highly transcribed putative extensin in heartwood/sapwood transition zone in black walnut.

28. Martínez-García PJ, Crepeau MW, Puiu D, Gonzalez-Ibeas D, Whalen J, et al. (2016) The walnut (juglans regia) genome sequence reveals diversity in genes coding for the biosynthesis of nonstructural polyphenols. The Plant Journal.

29. Chakraborty S, Britton M, Martínez-García P, Dandekar AM (2016) Deep RNA-seq profile reveals biodiversity, plant–microbe interactions and a large family of NBS-LRR resistance genes in walnut (juglans regia) tissues. AMB Express 6: 1.

30. Chakraborty S, Martínez-García PJ, Dandekar AM (2016) Yeatsam analysis of the walnut and chickpea transcriptome reveals key genes undetected by current annotation tools. F1000Research 5.

31. Chakraborty S (2017) Mcf-7 breast cancer cell line pacbio generated transcriptome has˜300 novel transcribed regions, un-annotated in both refseq and gencode, and absent in the liver, heart and brain transcriptomes. bioRxiv: 100974.

